# Circularization of genes and chromosome by CRISPR in human cells

**DOI:** 10.1101/304493

**Authors:** Henrik Devitt Møller, Lin Lin, Xi Xiang, Trine Skov Petersen, Jinrong Huang, Luhan Yang, Eigil Kjeldsen, Uffe Birk Jensen, Xiuqing Zhang, Xin Liu, Xun Xu, Jian Wang, Huanming Yang, George M. Church, Lars Bolund, Birgitte Regenberg, Yonglun Luo

## Abstract

Extrachromosomal circular DNA (eccDNA) and ring chromosomes are genetic alterations found in humans with genetic disorders and diseases such as cancer. However, there is a lack of genetic engineering tool to recapitulate these features. Here, we report the discovery that delivery of pairs of CRISPR/Cas9 guide RNAs into human cells generate functional eccDNAs and ring chromosomes. We generated a dual-fluorescence eccDNA biosensor system, which allows us to study CRISPR deletion, inversion, and circularization of genes inside cells. Analysis after CRISPR editing at intergenic and genic loci in human embryonic kidney 293T cells and human mammary fibroblasts reveal that CRISPR deleted DNA readily form eccDNA in human cells. DNA in sizes from a few hundred base pairs up to a 47.4 megabase-sized ring chromosome (chr18) can be circularized. Our discoveries advance and expand CRISPR-Cas9 technology applications for genetic engineering, modeling of human diseases, and chromosome engineering.

**One Sentence Summary:** CRISPR circularization of DNA offers new tools for studying eccDNA biogenesis, function, chromosome engineering, and synthetic biology.

---

The Clustered Regularly Interspaced Short Palindromic Repeats (CRISPR) and CRISPR-associated protein 9 (Cas9) is an adaptive immune system in bacteria and archaea that eliminate phages (1). Mediated by a small guide RNA (gRNA), the endonuclease Cas9 is harnessed for gene editing and has quickly become a highly attractive and powerful tool in basic and applied research (2–4). The fidelity and nuclease-precision of the CRISPR-Cas9 system insures no or minimal off-target effects and facilitates fast, functional gene studies of e.g. site-directed mutations in eukaryotic cells. Moreover, CRISPR-Cas9 mediated gene knockouts can be created though a pair of gRNAs that triggers dual double-stranded DNA breaks, e.g. at early or critical exons, leading to a DNA deletion and functionality disruption of the gene encoded protein (3, 4) or the inactivation of porcine endogenous retroviruses (5).

However, it is commonly thought that excised DNA after dual restriction by CRISPR-Cas9 will degrade or dilute out through cell proliferations, yet the faith of deleted chromosomal DNA is still unknown. Any nuclear double-stranded DNA breakage will in principle be recognized by the DNA repair machinery, which could result in non-homologous end joining or homology-directed repair of the deleted linear DNA fragment, leading to a DNA circle. DNA deletions and DNA inversions have previously been shown to form after dual restriction by CRISPR-Cas9 (6, 7) but it remains to be resolved whether deleted DNA fragments also form extrachromosomal circular DNA (eccDNA). Genome-scale studies have recently shown that eccDNAs are common elements in human cells (8, 9) and malignant tumors often carry oncogene amplifications on eccDNAs, known as double minutes (10–13). The mutational processes, leading to eccDNA, appear to occur mostly at random, although sub-sets of eccDNAs appears to form more frequently (8, 9, 14). A direct connection between DNA deletions and eccDNA formation has only been found sporadically (8, 12, 15, 16) and for 50 years, cancer studies of double minutes/eccDNA have been prohibited by limited eccDNA detection techniques, underlying the need for directed genetic tools to facilitate eccDNA analyses.

Here we describe such a tool and explore the induction of eccDNA formation and DNA deletions/insertions after dual DNA cuts by CRISPR-Cas9. To enable real-time monitoring of eccDNA biogenesis in cells, we designed and synthesized an eccDNA biosensor (ECC) with constitutive gene-expression: pCAG-ECC (Fig. S1) as well as a tetracycline inducible biosensor: pTRE-ECC (Fig. 1 and S2). The principle for each designed ECC-cassette was to mark the outcome after DNA repair of two introduced site-directed double-strand DNA breaks in cultured human cells, using CRISPR gRNAs (Cr1 + Cr2) (Fig. 1A and Fig. S3). A genomic DNA deletion, of the sequence encoding enhanced green fluorescent protein (Δ*EGFP*), would lead to functional *mCherry* expression after end joining with *mCherry* promoter (pCAG or pTRE). Likewise, *EGFP* expression was expected to mark circularization of the deleted DNA fragment, leading to an [*EGFP^circle^*]. Alternatively, an inversion could also lead to expression of both *EGFP* and *mCherry* but under a different expression scheme (Fig. 1A and Fig. S3). To experimentally test the dual-fluorescence ECC-system, we transiently transfected human embryonic kidney 293T cells (HEK293T) with pCAG-ECC and Cr1+Cr2, along with serial control transfections (Fig. S4 and S5). As expected, only cells transfected with pCAG-ECC and Cr1+Cr2 expressed both *EGFP* and *mCherry*. However, as transient transfection of the pCAG-ECC vector could not distinguish eccDNA from inversion events (Fig. S3) we tested the inducible pTRE-ECC in HEK293T. Here the majority of the cells, in absence of tetracycline, were EGFP positive, indicating [*EGFP^circles^*], while substantially less cells were mCherry positive (Fig. 1B-D, Fig. S6 and S7). Addition of tetracycline to cells enhanced both *EGFP* and *mCherry* expression, supporting formation of [*EGFP^circles^*], when targeted by CRISPR pairs, along with formation of deletions and inversions.

**Fig. 1.**
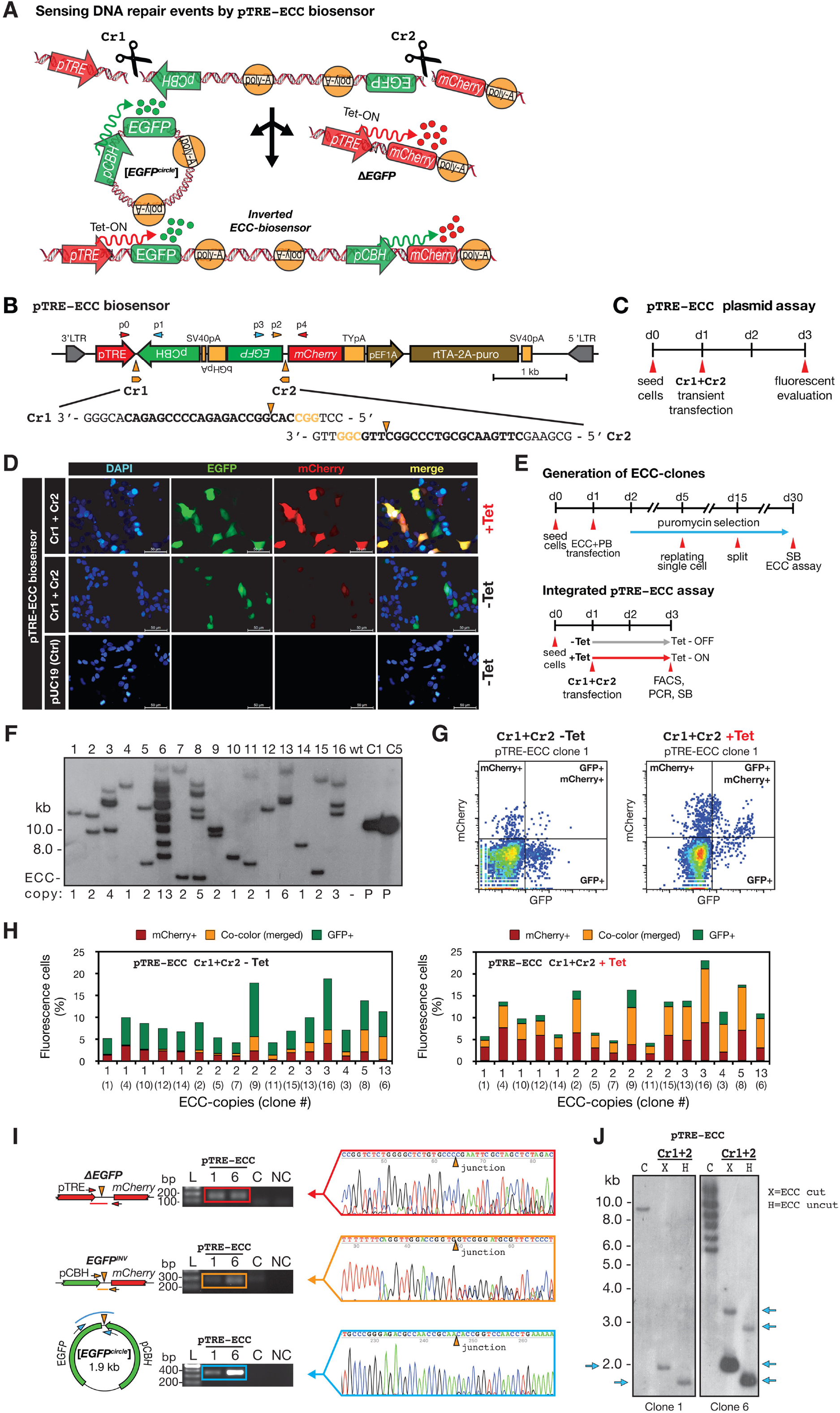
pTRE-ECC biosensor activation by CRISPR-pairs. (A) Model of pTRE-ECC biosensor after two double-stranded breaks by CRISPR-pairs (Cr1 and Cr2), leading to [*EGFP^circle^*] and Δ*EGFP* or *EGFP* inversion. (B) Scheme of pTRE-ECC biosensor. (C) Experimental outline. (D) pTRE-ECC plasmid assessment by fluorescence microscopy after Cr1+Cr2 using pUC19 as control (uncut). (E, upper part) Outline of stable genomic integration of pTRE-ECC in HEK293T. (F) Copy-number assessment by Southern blot with EGFP-probe after *KpnI* digestion. (E, lower part) Outline for Cr1+Cr2 activation of integrated pTRE-ECC. (G) Representative result from FACS analysis of clone 1 in the absence or presence of tetracycline (Tet). (H) Histograms of fluorescence cell percentages of all 15 isolated pTRE-ECC clones after Cr1+Cr2 and FACS analysis. (I) PCR and Sanger sequencing validation of genotypes, displayed in A, after Cr1+Cr2 for pTRE-ECC clone 1 and clone 4. C, negative CRISPR control; NC, non-template control. (J) Southern blot, probed with EGFP on HEK293T purified and digested DNA from untreated (C, *KpnI*) and Cr1+Cr2 treated cells. X=*XbaI*, H=*HindIII*.

Transient transfection is known to overload cells with many copies of plasmids. To overcome plasmid overloading and to distinguish inversions from eccDNAs, we established stable HEK293T biosensor reporter cells by the PiggyBac transposon system. Nine pCAG-ECC and 15 pTRE-ECC clones were established. Each clone carried from 1 up to 13 copies of the biosensor cassette, assessed by Southern blotting (Fig. 1E-G, Fig. S8 and S9). Fluorescence-activated cell sorting (FACS) analysis of all individual cell clones, induced by Cr1+Cr2, revealed correlation between the ECC-biosensor copy number per cell and median levels of EGFP and mCherry signals (*ρ*= 0.91 pCAG; *ρ*= 0.91 pTRE-Tet; *ρ*= 0.92 pTRE +Tet, Pearson rank test). It further validated formation of functional eccDNA after dual CRISPR-editing (Fig. 1H, Fig. S8 and S9). Actual formation of the 1.9 kb-sized [*EGFP^circle^*], along with *EGFP* deletion (Δ*EGFP*) and *mCherry* inversion, was confirmed by Sanger sequencing of PCR products and [*EGFP^circles^*] were further confirmed by Southern blot analysis for both the pTRE-and the pCAG-ECC system (Fig. 1I-J and Fig. S10).

Endogenous eccDNA has previously been found in both tumor and normal cells (8, 10). We next investigated whether dual CRISPR-targeting also generated eccDNA in tumor (HEK293T) and normal cells (human mammary fibroblasts, HMF). EccDNAs are commonly derived from intergenic regions and often consist of repeated sequences (8, 17). Thus, we designed two CRISPR gRNAs (Cr3+Cr4) targeting a site on chromosome 1 (chr1:159,708,525-159,709,943), which is located 2 kb downstream of the C-reactive protein encoding gene, *CRP*, and contains a SINE (AluYk4) element (Fig. 2A). We first evaluated the deletion efficiency to ~ 70 % after dual CRISPR-targeting (Cr3+Cr4) in HEK293T cells (Fig. 2B-C). We then confirmed CRISPR inversion events in these cells by PCR and Sanger sequencing (Fig. 2D). Finally, and as anticipated, Cr3+Cr4 treatment led to circularization of the deleted DNA, resulting in formation of the 0.57 kb-sized [*dsCRP^circle 1q23.2:0.57 kb^*] (Fig. 2E). Since HEK293T cells are polyploidy and a CRISPR duplication event potentially could give a false-positive PCR signal for eccDNA (Fig. S11), we optimized purification and exonuclease protocols (n=3) to remove all linear/chromosomal DNA (Fig. S12). The optimal cell lysate PCR protocol allowed us to detect [*dsCRP^circle 1q23.2:0.57 kb^*] in 10,000 cells, using outward directing oligos. Controls confirmed complete removal of linear DNA but not plasmid DNA (Fig. S12). EccDNA purification protocols were repeated in normal human mammary fibroblast (HMF) cells, reaching comparable results and conclusions (Fig. S13). Thus, the cell lysate method was hereafter used for all proceeding experiments (Fig. S12 and S13). Taken together, we found that dual CRISPR restriction, in close proximity, leads to DNA circularization of intergenic DNA in both normal and cancer cells, supporting our observations from CRISPR-pair activation of the ECC-biosensor system.

**Fig. 2.**
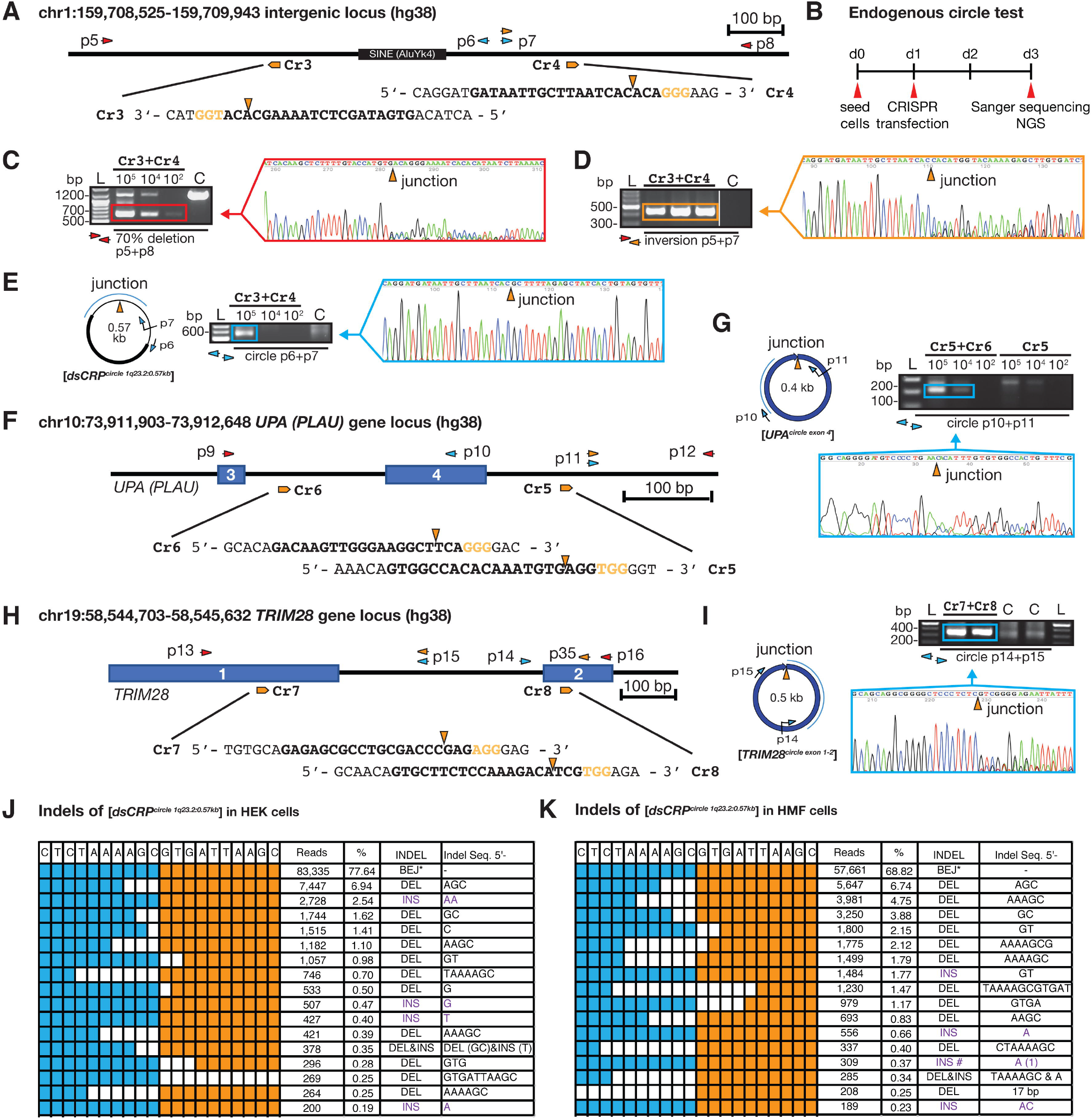
Endogenous DNA circularization by dual-CRISPR. (A) Chromosome 1 map 2 kb downstream of the *CRP* locus. gRNA; Cr3, Cr4; diagnostic oligos; p5 to p8. (B) Experimental outline. (C-E) Gel-images of corresponding PCR products across junctions after testing for (C) DNA deletion, 0.64 kb; wildtype, 1.23 kb), (D) inversion and (E) circular DNA, next to resultant chromatograms from Sanger sequencing. (F) Chromosome 10 map at the *PLAU (UPA)* locus. gRNA, Cr5 and Cr6, diagnostic oligos; p9 to p12. (G) PCR and sequencing confirmation of the [*<u>UPA</u>^circle exon 4^*]. (H) Chromosome 19 map at the *TRIM28* locus. (I) PCR and sequencing confirmation of the [*<u>TRIM28</u>^circle exon 1-2^*]. (J-K) Indel distribution across the junction of [*dsCRP^circle^*] generated in HEK293T and HMF cells, respectively.

Next, we tested induction of DNA circularization by CRISPR-pairs at two different genic loci in both normal and cancer cells as endogenous eccDNAs also are reported to derive from genic sequences (8, 10, 13, 18, 19). In line with findings of the [*dsCRP^circle 1q23.2:0.57 kb^*], Sanger sequencing of outward PCR products confirmed generation of the [*UPA^circle exon 4^*] and [TRIM28*^circle exon 1-2^*], leading to respective gene-disruption of the gene-encoding plasminogen activator (urokinase) and tripartite motif containing 28 (Fig. 2F-I and Fig. S14 and S15). Hence, endogenous eccDNA formation after dual CRISPRs seems to be a general outcome in cells, since it is cell-line independent and eccDNAs are formed at all positions tested.

Sanger sequencing of the circular junctions of [*EGFP^circle^*], [*dsCRP^circle 1q23.2:0.57 kb^*], [*UPA^circle exon 4^*] and [*TRIM28^circle exon 1-2^*], gave evidence of end-joining events. This supports the previous notion that the major DNA-repair pathway that appears to mediate eccDNA formation in mammalian cells is non-homologous end joining (NHEJ) (20, 21). To better understand the indel events, we further sequenced the DNA across the junction of the [*dsCRP^circle 1q23.2:0.57 kb^*] and the [*TRIM28^circle exon 1-2^*] by deep sequencing. For CRISPR-introduced *dsCRP* eccDNA, we observed 68.8% and 77.6% perfect blunt end-joining (BEJ) in HEK293T and HMF cells, respectively while the remaining 31.2% and 22.4% had a variable indel size up to maximum 17 bp (Fig. 2J-K). Similarly, the majority of [*TRIM28^circle exon 1-2^*] indels was less than 4 bp (79%) (Fig. S16). Though, for [*TRIM28^circle exon 1-2^*], we only observed 6.8% formed by BEJ while a majority (61%) had either one nucleotide insertion (43.3%) or deletion (13.2%) (Fig. S16). Such an indel distribution has previously been reported after CRISPR-editing (22, 23). In sum, the data indicates DNA repair of deleted DNA fragments from either NHEJ or microhomology-mediated end joining (MMEJ).

Succeeding experiments were carried out to evaluate the stability of *de novo* generated eccDNA after CRISPR DNA circularization, using the transgenic pTRE-ECC and pCAG-ECC clones with only one single copy insertion (Fig. 1F, clone 1 and Fig. S8B, clone 2, respectively). FACS experiments revealed dual expression of *EGFP* and *mCherry*, showing a rapid reduction in EGFP detectable fluorophores in solely green cells, propagated for more than one passage (1:3 cell-split ratio). In contrast, steady levels of red cells and co-color cells was observed in both pTRE-ECC and pCAG-ECC experiments (Fig. 3A-C and Fig. S17A-C). We interpreted co-colored cells with mCherry^high+^ and GFP^low+^ in pTRE-ECC (clone1, +Tet) as cells that likely contained Δ*EGFP* and [*EGFP^circles^*]. Over time, this group nearly vanished while the fraction of red and co-colored cells with low mCherry^low+^ and GFP^high+^ was steady, plausibly corresponding to deletion and *EGFP* inversion, respectively (Fig. 3C and Fig. S17C). When testing for presence of [*EGFP^circles^*] by outward PCR across the junction, we still found DNA products in all passages (Fig. 3D and Fig. S17D), supporting that sub-populations of cells contained [*EGFP^circles^*] for more than two weeks (6 passages). Hence, based on FACS analyses, the majority of *de novo* generated [*EGFP^circles^*] quickly became silenced and were gradually lost in cells over time. However, as [*EGFP^circles^*] were detected in all passages, degradation of circular DNA did not appear to be a rapid/active process.

**Fig. 3.**
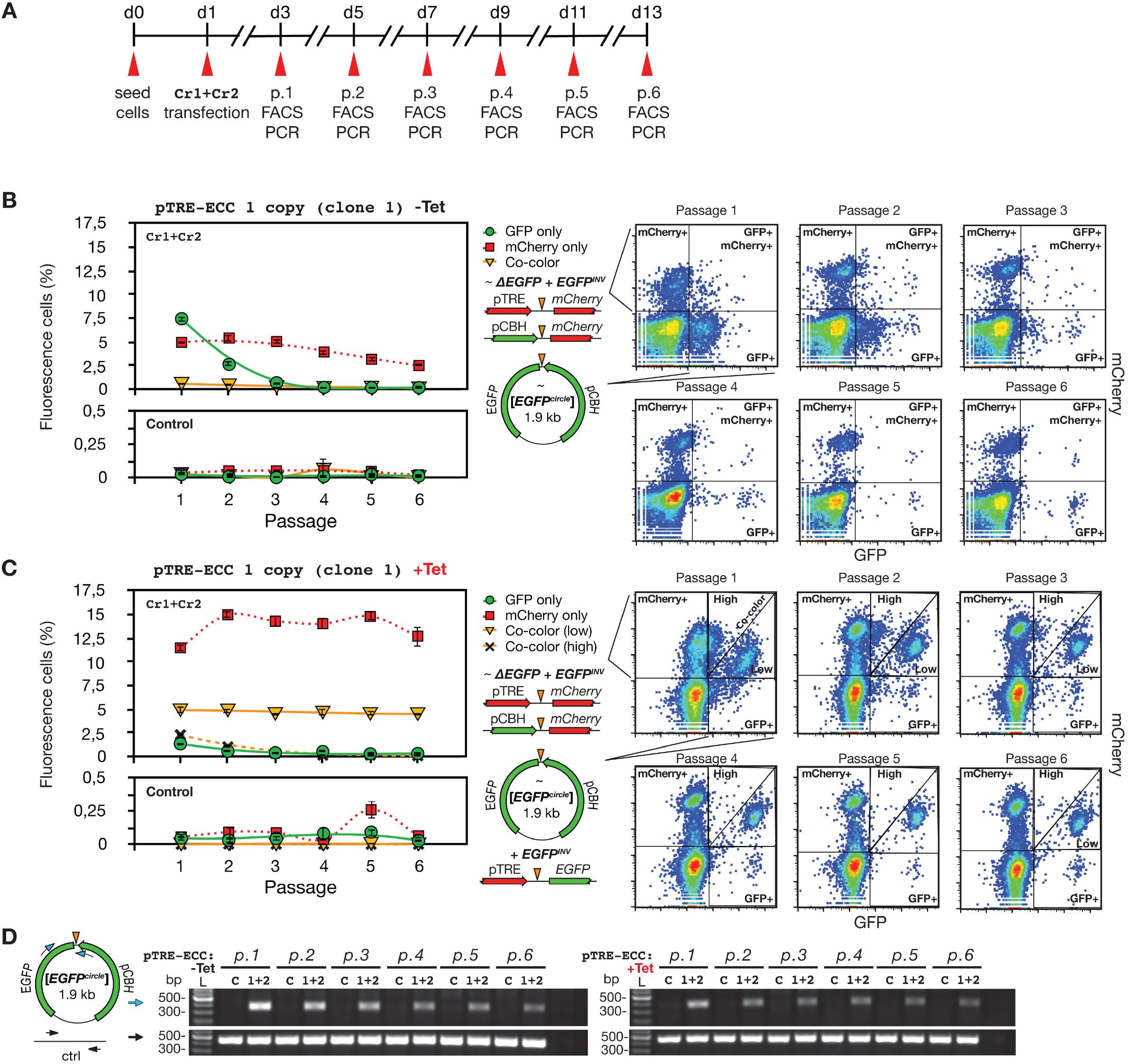
Time-course of [*EGFP^circle^*] expression and retention in cell culture. (A) Outline of time-course experiment. (B) Left, percent fluorescence cells from 1 to 6 cell passages (1:3 split-ratio) in absence of tetracycline (-Tet); right, corresponding FACS gating images. (C) As B description, cells propagated in +Tet conditions. (D) Outward PCR analysis of [*EGFP^circles^*] at passage 1 to 6; left,-Tet; right, +Tet. Cr1+Cr2 (1+2), CRISPR gRNAs; C, control - gRNA; ctrl, DNA template control.

Subsequently, we asked whether dual CRISPR-editing also could generate eccDNA in sizes larger than 2 kb. We generated targeted CRISPR-pairs to chromosome 1 and tested for eccDNA formation in sizes up to 207 kb. For all examined sites, we confirmed eccDNA formation after dual-cutting, while 1 cut controls did not led to eccDNA formation (Fig. 4A-C). Based on this data, double DNA breakage on the same chromosome seems to provide a high risk of DNA alterations in the form of DNA circularization and deletion.

**Fig. 4.**
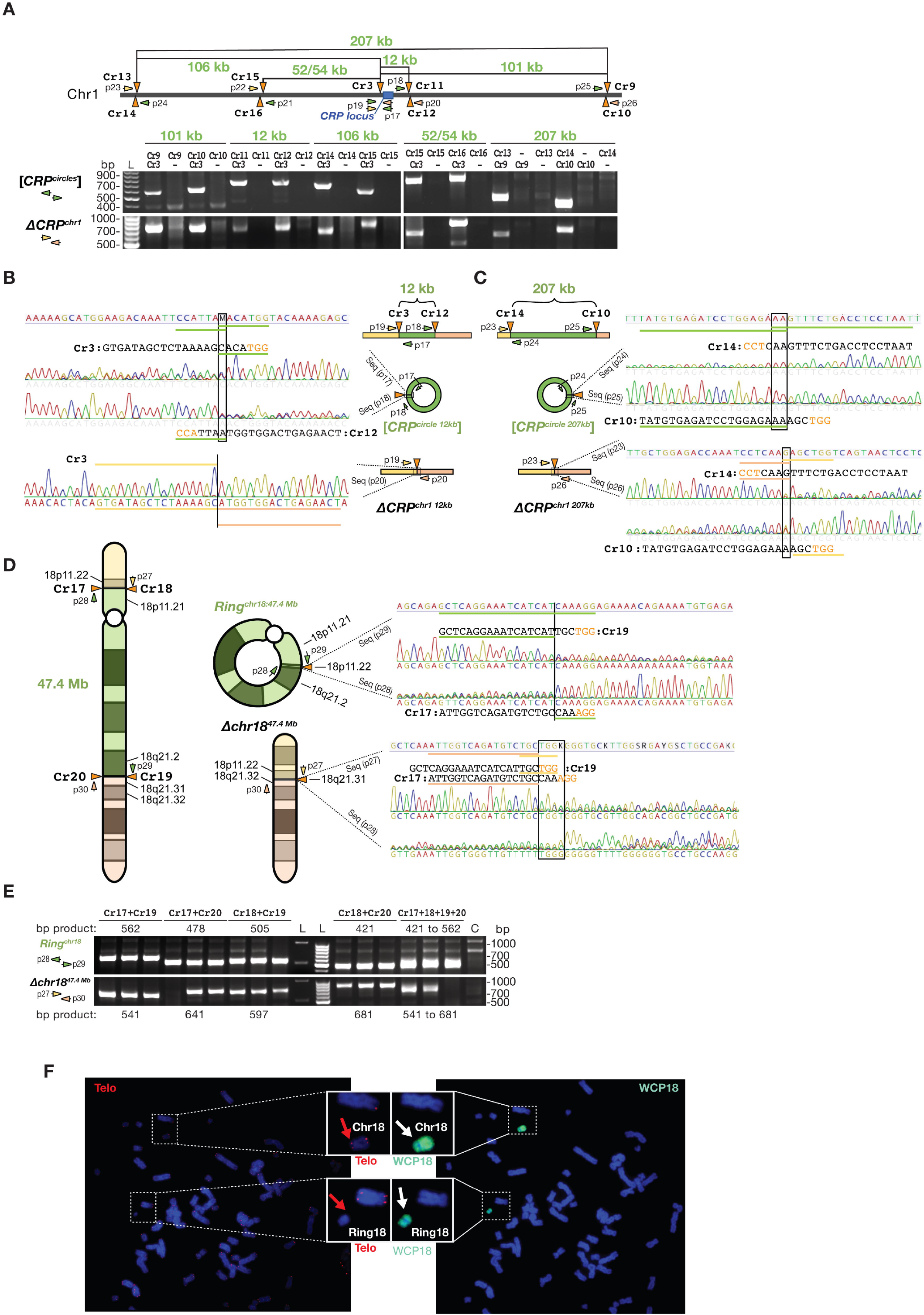
Circularization of kb and Mb-sized DNA fragments. (A, upper part) Schematic view of distances between targeted CRISPR gRNAs on chromosome 1 (Cr9 to Cr17) along with diagnostic oligos (p17 to p26) for PCR confirmation of formation of (A, lower part) kb-sized [*CRP^circles^*] and corresponding *CRP* deletions, Δ*CRP^chr1^*. (B-C) Chromatograms of Sanger sequencing across the junction of the [*CRP^circle 12 kb^*] and the *CRP^circle 207 kb^*] in both directions and chromatograms below confirming *CRP* deletions, Δ*CRP^chr1 12 kb^* and Δ*CRP^chr1 207 kb^*. (D) Left, schematic view of chromosome 18 with annotated Cr18 to Cr21 sites and resultant formation of *Ring^chr18:47.4 Mb^* and Δ*chr18^47.4Mb^* after dual CRISPR induction and DNA repair. Right, Sanger sequencing chromatograms confirming expected junctions. (E) Corresponding gel-images of amplified PCR products across junctions associated to D. Schemes, not drawn to scale. (F) Karyotyping, telomere staining and whole chromosome 18 painting of HEK293T cells treated with CRISPR-pairs. Blue, DPAI; Red, telemere; Green, Chromosome 18.

We next tested for CRISPR-induced circularization of chromosome 18, Ring^chr18^. Ring chromosomes are chromosomal alterations found in humans, which cause various developmental defects during fetal development and are associated to cancers (24). Currently, there is no tool for remodelling chromosome circularizations in cell culture. Ring formation has been reported for many chromosomes (24). Particular large deletions of chromosome 18, up to 36 Mb, are found at high frequency in 1 out of 40,000 births (25). When treating with CRISPR-pairs that targeted chromosome 18, we detected both efficient formation of large DNA deletions, Δchr18^47.4 Mb^ as well as Ring^chr18:47.4 Mb^. Sanger sequencing of amplified PCR products across the junction validated their structures (Fig. 4D-E). Karyotyping, telomere staining and whole chromosome 18 painting of HEK293T cells treated with CRISPR-pairs further confirmed the appearance of Ring^chr18:47.4 Mb^ with an efficiency of approximately 2% (2/113 metaphases analyzed, Fig. 4F), whereas no Ring^chr18:47.4 Mb^ was found in untreated cells (0/65 metaphases analyzed, Fig. S18). In addition, we observed that cells with the Ring^chr18:47.4 Mb^ genotype had lower fitness, as the Ring^chr18:47.4 Mb^ was eventually lost from cell culture when passaged multiple times. This result is not surprising as ring chromosomes are reported to be unstable (26).

## Conclusion

Here we report the discovery of precise CRISPR-mediated circularization of DNA in human cells. Upon simultaneous introduction of two double-stranded DNA breaks by CRISPR-pairs on the same chromosome eccDNA can form, likely mediated by the NHEJ/MMEJ DNA-repair pathways. Transient transfection of CRISPR-pairs can generate entire ring chromosomes inside cells and eccDNAs can be generated in a broad range of sizes from both genic and intergenic regions. We find that DNA circularization by CRISPR is cell-line independent and we consolidate previous studies (8, 11, 16, 18, 27), by showing that eccDNAs can be transcriptional active in cells, using a devised dual-fluorescence eccDNA biosensor tool.

Only few studies have previously looked at eccDNA stability over time (11, 19, 27). We established, from high-throughput FACS measurements, that eccDNA expression quickly becomes silenced in cells and eccDNAs are gradually lost over time. In addition, we find a propensity for eccDNAs to persist in sub-populations of cells after multiple passages, suggesting a rather slow decay of circular DNA. In yeast, ribosomal RNA genes are highly repetitive and known to frequently produce eccDNA known as [*rDNA^circles^*] (14) or ERCs (28). Accumulation of ERCs has been associated to cell senescence and aging of yeast (28) and potentially similar accumulation of eccDNAs in human cells could perhaps moderate the number of possible cell divisions, causing aging. Moreover, specific gene-encoding eccDNA were shown to: improve growth fitness in yeast (16); persist for more than 75 days in human cell culture (27); transmit drug-resistance vertically to progeny of crop weed (29).

Studies of chromosomal abnormalities in medical genetics, such as translocation, deletion, gene-fusion and ring chromosomes (30) can now be supplemented by applying dual CRISPR-technology. This method is especially useful for cancer research as around half of all tumors carry eccDNA with oncogenes (10, 11, 13). Using the dual CRISPR approach will improve our understanding of how eccDNA influence tumor biology and devised ECC-biosensor tools may be useful for genome synteny studies. It can further provide easier modeling of human genetic disorders, such as abnormalities caused by ring chromosomes (24, 26). In addition, the fact that ring chromosomes can be generated by CRISPR, provide valuable knowledge and novel concept arts to the design of synthetic chromosomes (31) and synthetic genomes (32).

## Acknowledgments

H.D.M. and B.R are supported by research grants from the Danish Council for Independent Research (FNU 6108-00171B). H.D.M. is also supported by the Carlsberg foundation (CF17-0226). L.L. is supported by the Lundbeck Foundation (R219-2016-1375). Xi. X. is supported by the China Scholarship Council. Y.L. is supported by the Danish Research Council for Independent Research (DFF-1337-00128), the Sapere Aude Young Research Talent Prize (DFF-1335-00763A), the Innovation Fund Denmark (BrainStem), the Lundbeck Foundation (R173-2014-1105) and Aarhus University Strategic Grant (AU-iCRISPR). We thank Claus Nielsen for helping with preparing the cells for karyotyping analysis and the technical help from our FACS CORE facility.

## Author contributions

H. D.M., L.L., Xi.X., B.R. and Y.L. designed experiments. H.D.M., L.L., Xi.X., T.S.P, Y.L. performed main experiments. H.D.M. and Y.L. finalized figures. H.D.M., L.L., Xi.X., Y.L., J.H., L.Y., E.K., U.B.J., X.Z., X L., X.Xu., J.W., H.Y. further analysed the data. H.D.M. and Y.L. wrote the manuscript with feedback from all authors.

## Supplementary Materials

Materials and Methods

Figures S1-S18

Tables S1-S2

References (1-**31**)

## References

1. R. Jansen, J. D. A. V. Embden, W. Gaastra, L. M. Schouls, Identification of genes that are associated with DNA repeats in prokaryotes. Mol Microbiol. 43, 1565–1575 (2002).

2. M. Jinek et al., A programmable dual-RNA-guided DNA endonuclease in adaptive bacterial immunity. Science. 337, 816–821 (2012).

3. L. Cong et al., Multiplex genome engineering using CRISPR/Cas systems. Science. 339, 819–823 (2013).

4. P. Mali et al., RNA-guided human genome engineering via Cas9. Science. 339, 823–826 (2013).

5. D. Niu et al., Inactivation of porcine endogenous retrovirus in pigs using CRISPR-Cas9. Science. 357, 1303–1307 (2017).

6. M. C. Canver et al., Characterization of genomic deletion efficiency mediated by clustered regularly interspaced short palindromic repeats (CRISPR)/Cas9 nuclease system in mammalian cells. J. Biol. Chem. 289, 21312–21324 (2014).

7. S. M. Byrne, L. Ortiz, P. Mali, J. Aach, G. M. Church, Multi-kilobase homozygous targeted gene replacement in human induced pluripotent stem cells. Nucleic Acids Research. 43, e21–e21 (2014).

8. H. D. Møller et al., Circular DNA elements of chromosomal origin are common in healthy human somatic tissue. Nat. Commun. 9, 1–12 (2018).

9. Y. Shibata et al., Extrachromosomal microDNAs and chromosomal microdeletions in normal tissues. Science. 336, 82–86 (2012).

10. K. M. Turner et al., Extrachromosomal oncogene amplification drives tumour evolution and genetic heterogeneity. Nature. 543, 122–125 (2017).

11. D. D. Von Hoff et al., Elimination of extrachromosomally amplified MYCgenes from human tumor cells reduces their tumorigenicity. Proceedings of the National Academy of Sciences. 89, 8165–8169 (1992).

12. S. M. Carroll et al., Double minute chromosomes can be produced from precursors derived from a chromosomal deletion. Molecular and Cellular Biology. 8, 1525–1533 (1988).

13. C. T. Storlazzi et al., Gene amplification as double minutes or homogeneously staining regions in solid tumors: Origin and structure. Genome Res. 20, 1198–12 (2010).

14. H. D. Møller, L. Parsons, T. S. Jørgensen, D. Botstein, B. Regenberg, Extrachromosomal circular DNA is common in yeast. Proceedings of the National Academy of Sciences. 112, E3114–E3122 (2015).

15. K. Durkin et al., Serial translocation by means of circular intermediates underlies colour sidedness in cattle. Nature. 482, 81–84 (2012).

16. D. Gresham et al., Adaptation to diverse nitrogen-limited environments by deletion or extrachromosomal element formation of the GAP1 locus. Proceedings of the National Academy of Sciences. 107, 18551–18556 (2010).

17. S. Cohen, N. Agmon, O. Sobol, D. Segal, Extrachromosomal circles of satellite repeats and 5S ribosomal DNA in human cells. Mobile DNA. 1, 11 (2010).

18. D. A. Nathanson et al., Targeted therapy resistance mediated by dynamic regulation of extrachromosomal mutant EGFR DNA. Science. 343, 72–76 (2014).

19. N. Vogt et al, Amplicon rearrangements during the extrachromosomal and intrachromosomal amplification process in a glioma. Nucleic Acids Research. 42, 13194–13205 (2014).

20. N. van Loon, D. Miller, J. P. Murnane, Formation of extrachromosomal circular DNA in HeLa cells by nonhomologous recombination. Nucleic Acids Research. 22, 2447–2452 (1994).

21. X. Meng et al., Novel role for non-homologous end joining in the formation of double minutes in methotrexate-resistant colon cancer cells. J. Med. Genet. 52, 135–144 (2015).

22. T. J. Cradick, E. J. Fine, C. J. Antico, G. Bao, CRISPR/Cas9 systems targeting β-globin and *CCR5* genes have substantial off-target activity. Nucleic Acids Research. 41, 9584–9592 (2013).

23. L. A. Lonowski et al., Genome editing using FACS enrichment of nuclease-expressing cells and indel detection by amplicon analysis. Nat Protoc. 12, 581–603 (2017).

24. R. S. Guilherme et al., Mechanisms of ring chromosome formation, ring instability and clinical consequences. BMC Med. Genet. 12, 171 (2011).

25. J. D. Cody et al., Congenital anomalies and anthropometry of 42 individuals with deletions of chromosome 18q. Am. J. Med. Genet. 85, 455–462 (1999).

26. C. P. Sodré et al., Ring chromosome instability evaluation in six patients with autosomal rings. Genet. Mol. Res. 9, 134–143 (2010).

27. G. Pauletti, E. Lai, G. Attardi, Early appearance and long-term persistence of the submicroscopic extrachromosomal elements (amplisomes) containing the amplified *DHFR* genes in human cell lines. Proceedings of the National Academy of Sciences. 87, 2955–2959 (1990).

28. D. A. Sinclair, L. Guarente, Extrachromosomal rDNA circles—a cause of aging in yeast. Cell. 91, 1033–1042 (1997).

29. D.-H. Koo et al., Extrachromosomal circular DNA-based amplification and transmission of herbicide resistance in crop weedAmaranthus palmeri. Proc Natl Acad Sci USA, 201719354 (2018).

30. D. Maddalo et al., In vivo engineering of oncogenic chromosomal rearrangements with the CRISPR/Cas9 system. Nature. 516, 423–427 (2014).

31. R. Chari, G. M. Church, Beyond editing to writing large genomes. Nat Rev Genet. 18, 749–760 (2017).

32. S. M. Richardson et al., Design of a synthetic yeast genome. Science. 355, 1040–1044 (2017).

